# Niche hypervolume analysis revealed that water pollution reduces reproductive success of endangered landlocked salmon but does not directly impact their population density

**DOI:** 10.1101/2022.03.21.485106

**Authors:** Yi-An Chung, Mark Liu, Tzu-Man Hung, Lin-Yan Liao, Sheng-Feng Shen

**Affiliations:** Biodiversity Research Center, Academia Sinica, Taipei, 11529 Taiwan; International Degree Program in Climate Change and Sustainable Development, College of Science, National Taiwan University, Taipei, 115, Taiwan; Institute of Ecology and Evolutionary Biology, National Taiwan University, Taipei, 10617 Taiwan; Wuling Station, Shei-Pa National Park, Taichung, Taiwan

**Author notes:** Corresponding author: Sheng-Feng Shen. Both authors contributed equally to this work (Yi-An Chung, Mark Liu).

## Abstract

The critical information for conserving endangered species is to identify how different niche dimensions affect key life history stages of a population. However, it is often difficult to quantify how each ecological niche dimension affects different life history stages because environmental factors may affect each fitness component of organisms to various degrees. Here, we applied the recently developed hypervolume method that follows the idea of Hutchinson’s n-dimensional hypervolume. We analyzed the niche space of different life history stages of the endangered landlocked salmon *Oncorhynchus masou formosanus*, the most southerly distributed of all salmonoids. We found no direct effect of water pollution on salmon population density but a significant negative effect on salmon reproductive success. Surprisingly, the niche hypervolume analysis showed that the size of hatching success niche hypervolume was only 17.9% and 18.4% of the natural redd site density or population density, respectively. This result demonstrates a much higher environmental requirement for salmon during the egg stage than that affecting population density or nesting environment. Our results suggest that understanding the behavioral and physiological mechanisms that influence crucial life history stages in the wild is critical to developing effective conservation programs, and the niche hypervolume is a valuable method to achieve this.

## Introduction

To understand the degree of threat and vulnerability of an endangered species, it is crucial to identify the key niche space of the species and the corresponding physical environment [1, 2]. Most studies have used observational methods to establish the correlation between occurrence and environmental factors to infer the niche space of focal species [3]. However, this approach does not help us understand the environmental impacts on the most critical life history stages, which have the most significant impact on the population [4, 5]. Moreover, a local population may be a sink population in a source-sink dynamic, so even if the population size is large, it does not necessarily mean that the environmental conditions at that site are the most suitable niche space for that population [4]. In other words, different fitness components and population density are not always positively correlated, and environmental conditions favorable for individual survival and reproduction may not be the same [6, 7]. Therefore, there have been calls to establish a mechanistic understanding of species distribution, i.e., to understand the causal relationships of how environmental factors affect the fitness components of organisms (e.g., reproduction or survival), to truly understand the ecological niches that are critical to populations and to develop conservation strategies for endangered species accordingly [5].

The Formosan salmon (*Oncorhynchus masou formosanus*) is a landlocked salmon that exists only in the upper reaches of the Tajia River in Taiwan and is also the lowest latitude salmonid population [8]. Since the 1970s, the population of Formosan salmon has declined dramatically due to agricultural land development, hunting, and habitat fragmentation caused by check dams [9]. Originally widely distributed in the upper watershed of the Tajia River, only 200 individuals remain in a 5-km-long stretch of the Chichiawan Creek, Taiwan [10]. Therefore, the Formosan salmon was listed as an endangered species by the Taiwan government in 1989 and by IUCN in 1996, and Chichiawan Creek was designated as a wildlife refuge in 1997. Thanks to the efforts of the government, researchers, and conservationists, the population of Formosan salmon has increased to tens of thousands of fish. In addition, many studies investigate the reproductive biology of Formosan salmon [11] and the relationship between population size and the environment [12]. However, no systematic studies have been conducted on the reproductive ecology of wild Formosan salmon populations in Taiwan. Here we apply the concept and analysis of niche hypervolume [13, 14] to quantify the niche spaces of Formosan salmon during survival (population size), and during breeding, including redd sites and hatching rate. We hope to understand the most critical environmental factors that affect this endangered species by quantifying the niche space of different life history stages. We believe that this study will allow us to conserve the Formosan salmon more effectively and provide other studies on how the concept of niche hypervolume can be used to help conserve endangered species.

## Method

### Study areas

During the breeding season from late September to late November 2021, we inspected every transect at a frequency of 5-6 days to locate and record redd sites of Formosan salmon. We selected six study transects in Wuling area (figure S1). Transect 1 is located between Dam No. 4 and Dam No. 5 in Taoshan West Creek, with a study reach length of 600 m and an elevation of 1900 m. Transect 2 is located in the upper reaches of Chichiawan Creek between check dams No. 2 and No. 3, with a study reach length of 870 m. Transect 3 is located in the middle reaches of Chichiawan Creek, downstream of Check Dam No. 2, with a study length of 1100 m. Agricultural land on the hillside is mainly cultivated for fruits and large recreational purposes campsites in this reach. Transect 4 is upstream of check Dam No. 1, where the length of the Creek investigated is 380 meters. Transect 5 is located downstream of the confluence of Chichiawan Creek with Khaoshan Creek, and the length of the studied Creek segment is 370 m. Transect 6 is located downstream of the confluence of Chichiawan Creek and Yousheng Creek, with a length of 520 meters. Two transects were selected for the study in Hehuan Creek (figure. S2). Transect 1 is located at an elevation of 2,550 m above sea level in the upper valley of Hehuan Creek, and the vegetation along the creek is mostly undisturbed primary forest. Transect 2 is located in the lower valley of Hehuan Creek at an elevation of 2,250 meters. The vegetation along the creek comprises primary forest and vegetable gardens cultivated for agriculture. The length of this study section is 980 meters.

### Natural redd sites and microhabitat measurements

Formosan salmon females use their tails to excavate the riverbed and lay their eggs in the gravel gaps during the breeding season. The males drive other males away, perform courtship behaviors on the females, and then mate with them. Therefore, we identified redd sites by observing the behavior of salmon and the characteristics of salmon redd sites. After redd sites were recorded, we measured pH, dissolved oxygen (mg/L), and conductivity (mS/cm) at redd sites with a multi-parameter water quality meter (U-51; HORIBA), and flow velocity with a flow meter (438-115; HYDRO-BIOS). The water quality and velocity were measured at 40 randomly selected locations in each of the Chichiawan Creek and the Hehuan Creek as a control group. We divided the study reaches of Chichiawan Creek and Hehuan Creek into 1364 and 1119 cells on the map and then used R to randomly select the cell numbers as random locations to compare with the actual redd locations. We used this method to determine the redd site preference of Formosan salmon. We used an ibutton (DS1925; The iButton® Maxim Integrated) to measure water temperature, and the temperature was recorded every 30 minutes for the entire study period. We set up 27 and 24 water temperature stations in Chichiawan Creek and Hehuan Creek, respectively. We analyzed the effects of average water temperature, diurnal temperature range, daily maximum temperature, and daily minimum temperature on the reproductive performance of Formosan salmon. Each temperature variable was averaged over 24 hours per day and then averaged over each day of the sampling period. We also used water quality and water temperature data from randomly selected points in the surveyed creeks to analyze the effects of water quality and water temperature on salmon population density.

### Hatching rate

We conducted artificial hatching experiments to study the relationship between the hatching rate and the environment after the natural reproduction of salmon in Taiwan has ended to avoid interfering with natural reproduction. At the end of November, we selected 17 and 15 sampling sites in the Chichiawan and Hehuan Creeks, respectively, and set up three incubation boxes (Whitlock Vibert box) [15] at each site. The eggs came from a captive colony at the Formosan salmon Ecology Center in Shei-Pa National Park. We followed the Whitlock Vibert box instructions to construct suitable breeding sites in the riverbed where the salmon were nesting, using tools such as shovels. We set up the incubation box in a hole in the riverbed at a depth of about 10-20 centimeters and covered it with gravel after placing it in the box to ensure that it was upright and not exposed to sunlight. Three weeks after burying the Whitlock Vibert boxes, we removed the boxes from the streambed and counted the number of eggs, dead eggs, alevins, and dead alevins. We measured water depth, velocity, and water quality once after setting up the boxes and before removing them. Water temperature was measured by ibutton at 30-minute intervals throughout the experiment.

### Population Density

Since Formosan salmon is an endangered species, volunteers, staff, and researchers organized by Shei-Pa National Park, have used snorkel surveys to quantify the relative population density once or twice a year in each creek since 1987 [16]. We surveyed salmon populations in Chichiawan Creek and Hehuan Creek separately for 2 days each during the summer of 2021. The survey was conducted from 10:00 to 17:00 each day, with two counters in each reach simultaneously snorkeling upstream from both sides of the creek and counting the number of fish along the creek bank [16].

### Data Analysis

We use the GLM model with binomial distribution to compare water quality, velocity, and temperature data between natural redd sites and randomly selected control sites. We used the sampling creek ID as the random effect and water velocity, water quality, and water temperature as the fixed effect. We also modeled the hatching rate and the water quality, velocity, and water temperature sampled in the early and late periods separately in the LMM. We compared the AIC indices of the different models and examined the significance of the factors by using the sampled creek ID as a random effect. We compared the difference in water depth and velocity between the early and late sampling periods using a t-test right tail test. The relationship between population density, redd density, and hatching rate was examined by correlation analysis. To compute the alpha diversity between the hypervolumes of environmental niches, we used the alpha function in the “BAT” package (version 2.7.1) [20-21]. All data analysis was completed using R version 4.1.2 [17].

## Results

To quantify the environmental effects on Formosan salmon survival and reproduction, we selected Chichiawan Creek and Hehuan Creek as study sites. The reproductive performance included quantifying natural redd site selection and hatching rates using artificial Whitlock Vibert Boxes. Individual survival was represented by relative density surveys. To understand the impacts of the environment on the Formosan salmon population, we measured water quality (n=80, 40 sites in each creek) and recorded water temperature (n=50, n=27 in Chichiawan Creek, and n=23 in Hehuan Creek) in randomly-selected sites in different creeks. We found that water quality affected the relative population density of Formosan salmons. When the pH was closer to neutral, the population density was higher (table S1).

For reproduction, we tested redd site selection in Formosan salmon by comparing differences in water quality, flow velocity, and water temperature between natural redd sites (n=136 and n=96 in Chichiawan Creek and Hehuan Creek, respectively; figure 1a and 1b) and randomly selected sites in each creek (n=80, 40 sites in each creek, Table.1). We found that Formosan salmon preferred to build redd sites in locations with pH close to neutral (figure 1c), lower conductivity, and slower flow velocity (figure 1d; table S2).

**Figure 1.**
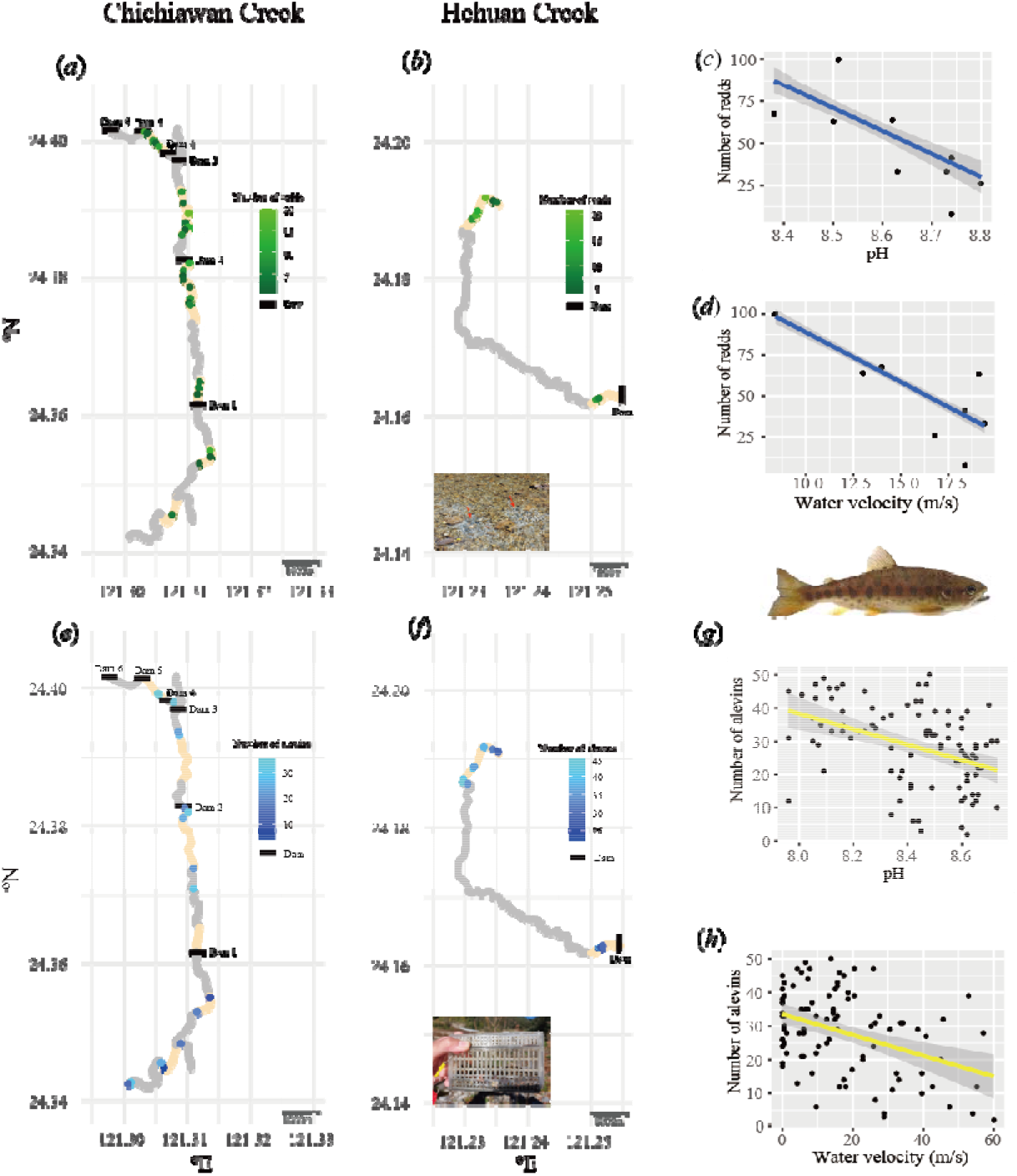
The distribution of redd density in Chichiawan Creek (a) and Hehuan Creek (b) and the relationship between redd density and water pH (c) and flow velocity (d). Distribution of the number of alevins hatched in the Whitlock Vibert box experiments in Chichiawan Creek (e) and Hehuan Creek (f) and the relationship between the number of alevins hatched and the water pH (g) and flow velocity (h).

To quantify the hatching rate, we selected 32 sample sites in Chichiawan Creek (n=17) and Hehuan Creek (n=15) close to natural redd sites. Three Whitlock Vibert boxes were placed at each site, each box containing 50 eyed eggs. The hatchlings ranged from 2-44 fish per box were retrieved after the experiment (figure 1e and 1f). We measured the environmental factors twice at the beginning and before the end of the experimental setup, respectively. The environmental factors we measured include pH, conductivity, dissolved oxygen, flow velocity, depth, average daily temperature (Tmean), average daily maximum temperature (Tmax), average daily minimum temperature (Tmin), and average DTR. We also found that the water level was higher in the first measurement than in the second (figure S2, t-test, p<0.001), so we analyzed the environmental factors for each of the two measurements. For the first measurement, we found that the factors that best explain the effect of hatching rate in the case of high-water level are the flow velocity, pH, DTR, and the interaction between DTR and flow velocity (AIC=707.34, table S3). All of these factors had significant effects on the hatching rate in the linear mixed model (LMM analysis; table S4). Except for the interaction between DTR and flow rate, all other factors had a negative effect on the hatching rate. That is, the closer the pH to neutral (figure 1g) and the lower the DTR, the higher the hatching rate. On the other hand, we found that flow velocity had a negative effect on hatching rate when DTR was low and a positive effect on hatching rate when DTR was high (figure 1h). While analyzing the results of the second measurement, we found that the factors that best explain the hatching rate were dissolved oxygen, conductivity, Tmin, the interaction between Tmin and conductivity (AIC=692.08, table S5), but only dissolved oxygen and Tmin significantly affect them in the LMM analysis (table S6). When the dissolved oxygen, conductivity, and Tmin were lower, the hatching rate was higher.

In addition, we explored the relationship between population density, redd site density, and hatching rate of Formosan salmon. We found that population density was positively correlated with redd site density (r=0.71, p=0.047). Redd site density was also highly positively correlated with hatching rate (r=0.54, p<0.001), but population density was not correlated with hatching rate (r=0.33, p=0.002). Finally, we used the niche hypervolume method to quantify the size of the niche hypervolume of Formosan salmon during the survival and reproductive stages. We included pH, water flow velocity, dissolved oxygen (DO), and water conductivity, which are four environmental factors that have significant effects on salmon survival or reproduction, to calculate the size of niche hypervolume. We found that the size of the niche hypervolume was similar for the population density and natural redd sites of Formosan salmon, but the niche hypervolume for reproductive success represented by hatching rate (0.177; figure 2a) was only 17.9% and 18.4% of the natural redd density (0.990; figure 2b) and population density (0.962; figure 2c) and, respectively (Figure 2d).

**Figure 2.**
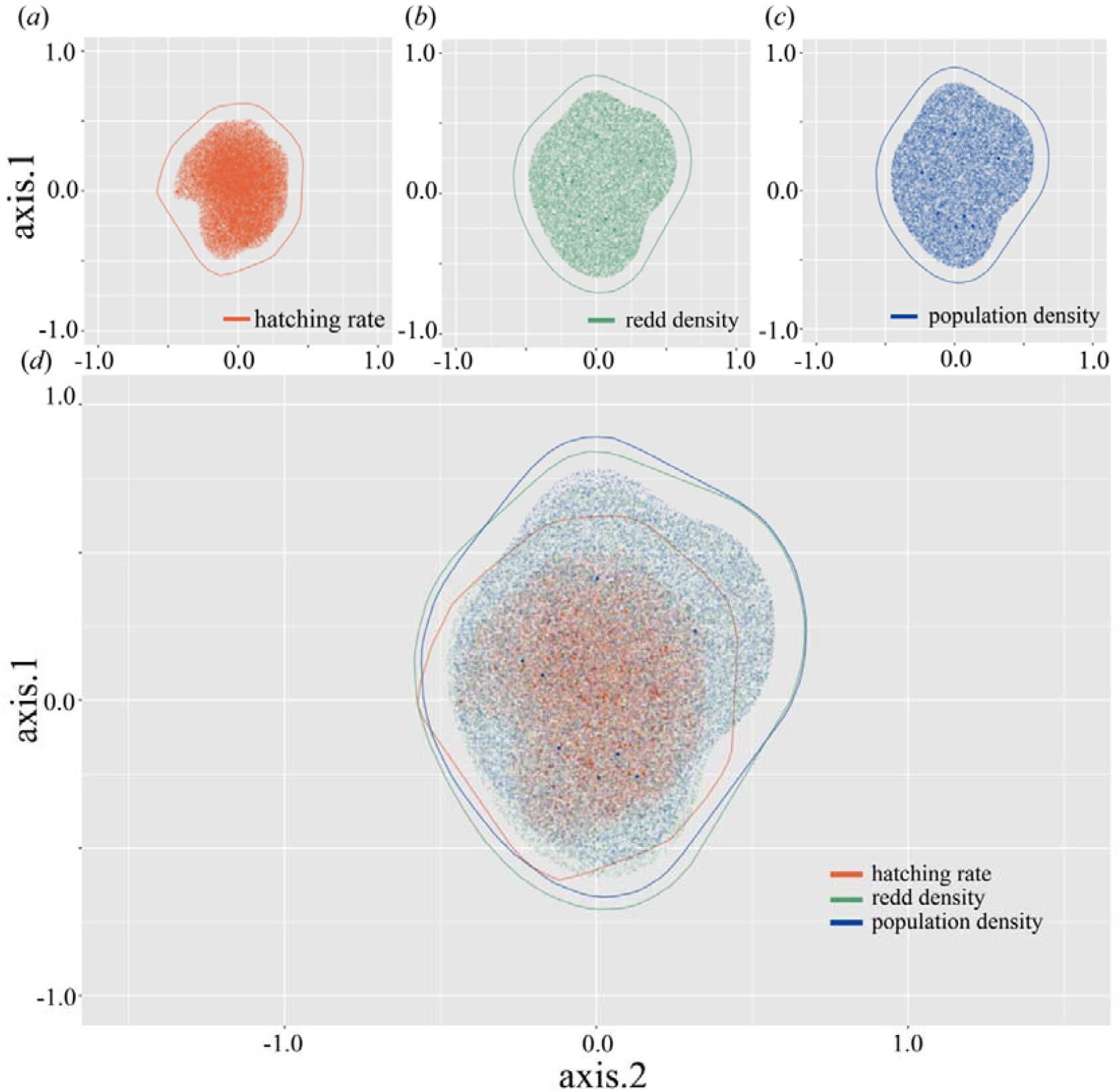
The niche hypervolume of hatching rate (a), redd density (b), and population density (c) and a comparison of the three (d). Key environmental factors that affect salmon reproduction or survival, including water pH, flow velocity, dissolved oxygen (DO), and conductivity, were used to calculate the size of the niche hypervolume.

## Discussion

We used the niche hypervolume method to quantify the size of the niche space for the environmental factors most important to the survival and reproduction of Formosan salmon. We found that the niche hypervolume for successful breeding is the smallest and therefore indicates the highest environmental requirement for successful reproduction of Formosan salmon. In other words, although water quality does not have a significant effect on the relative density of the Formosan salmon population, water quality has a substantial effect on their redd site selection and reproductive success. In addition, the results of our artificial breeding test by Whitlock Vibert Box on reproductive performance also showed that pollution had a more negligible effect on reproduction during periods of high-water level. In contrast, water pollution had a very significant negative impact on breeding performance during periods of low water level. Therefore, from a conservation perspective, management could use our results to focus on improving water quality during the breeding season, such as reducing the number of overnight visitors allowed during the breeding season, to increase the reproductive success of Formosan salmon.

More broadly, with advances in niche hypervolume analysis methods, we now have a better tool to analyze the size of niche hypervolumes for endangered species at different life history stages. Having used the concept of hypervolume to quantify niche space, we can also apply the concepts of ecological niche space (abstract environmental factors) and biotope (the actual physical environment) to quantify the availability of suitable habitat for endangered species at crucial life history stages [1]. Previous studies have used species distribution models to quantify differences in environmental requirements between breeding and non-breeding populations[18]. However, the most significant advantage of using the hypervolume approach is that the size of niche space - an environmental factor critical to the organism - can be compared across life history stages [19]. This will allow us to quantify further the size of biotope (actual habitat) suitable for a focal species based on identified vital niche dimensions such as river flow velocity and pH. Thus, our study echoes previous calls for the development of mechanistic species distribution models [3-5]. Ultimately, we believe that, especially for endangered species, we need to understand not only the environmental factors that influence presence/absence or population size but also the behavioral and physiological mechanisms that influence crucial life history stages in the wild in order to fully understand the mechanisms that influence the animal distribution and carry out effective conservation programs.

## Data accessibility

The data are provided in the electronic supplementary material

## Authors’ contributions

Y.-A. C., M. L., L.-Y. L. and S.-F.S. designed the study. Y.-A. C. and M. L. performed the field study and collected breeding data. L.-Y. L. collected population density data. T.-M. H. and Y.-A. C. performed the hypervolume analysis. Y.-A. C. analyzed the data. S.-F. S. conceived the project. Y.-A. C., M. L., and S.-F.S. wrote the first draft of the paper. All authors reviewed, edited the paper, and gave final approval for publication.

## Acknowledgments

We thank Alven Liao, William Yan-Lun Shen, and the staff of Shei-Pa National Park for their help with fieldwork and logistics.

## Supplementary material

**Figure S1.**
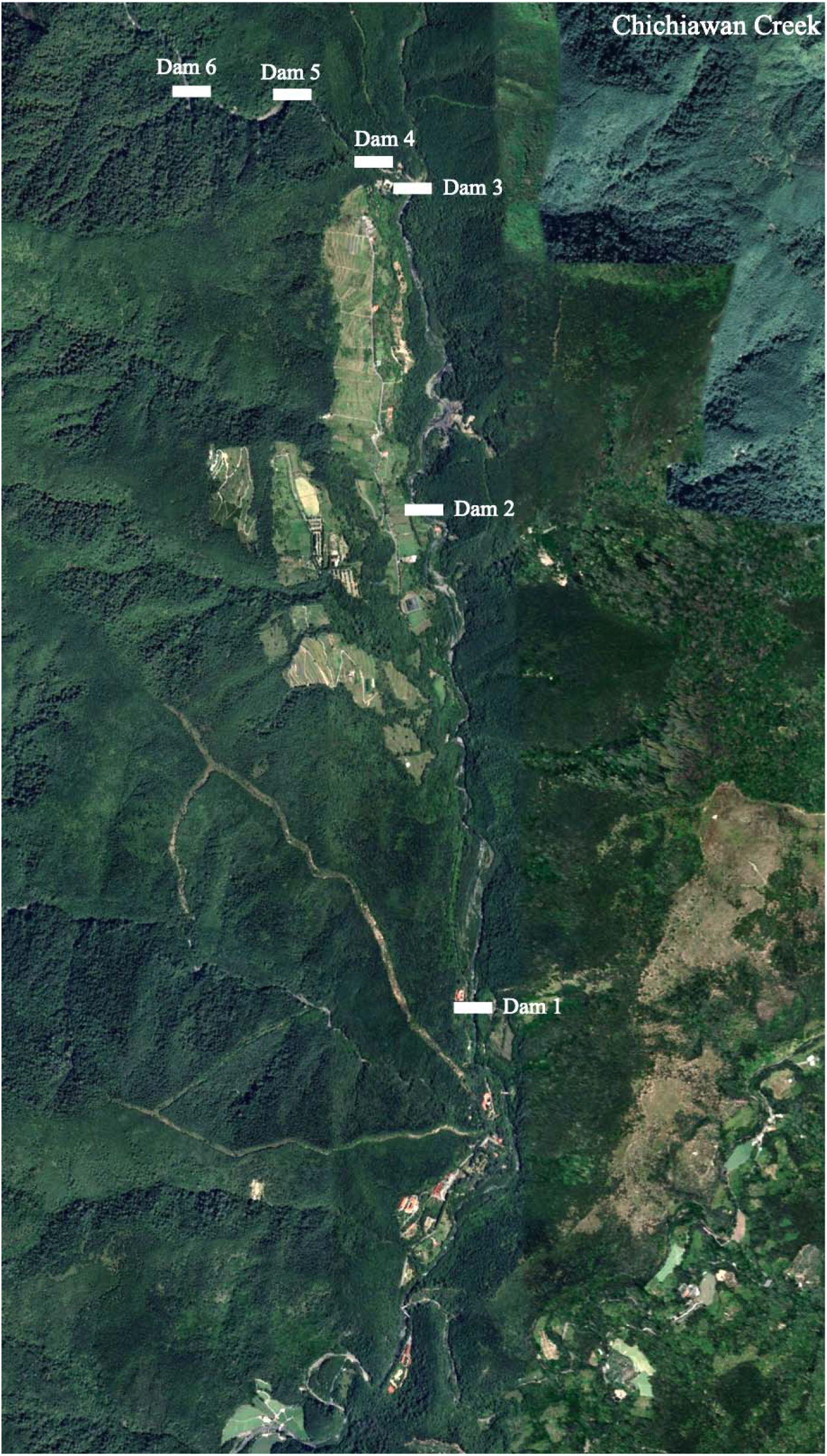
A satellite photo of the study area of Chichiawan Creek. Image from Google Earth.

**Figure S2.**
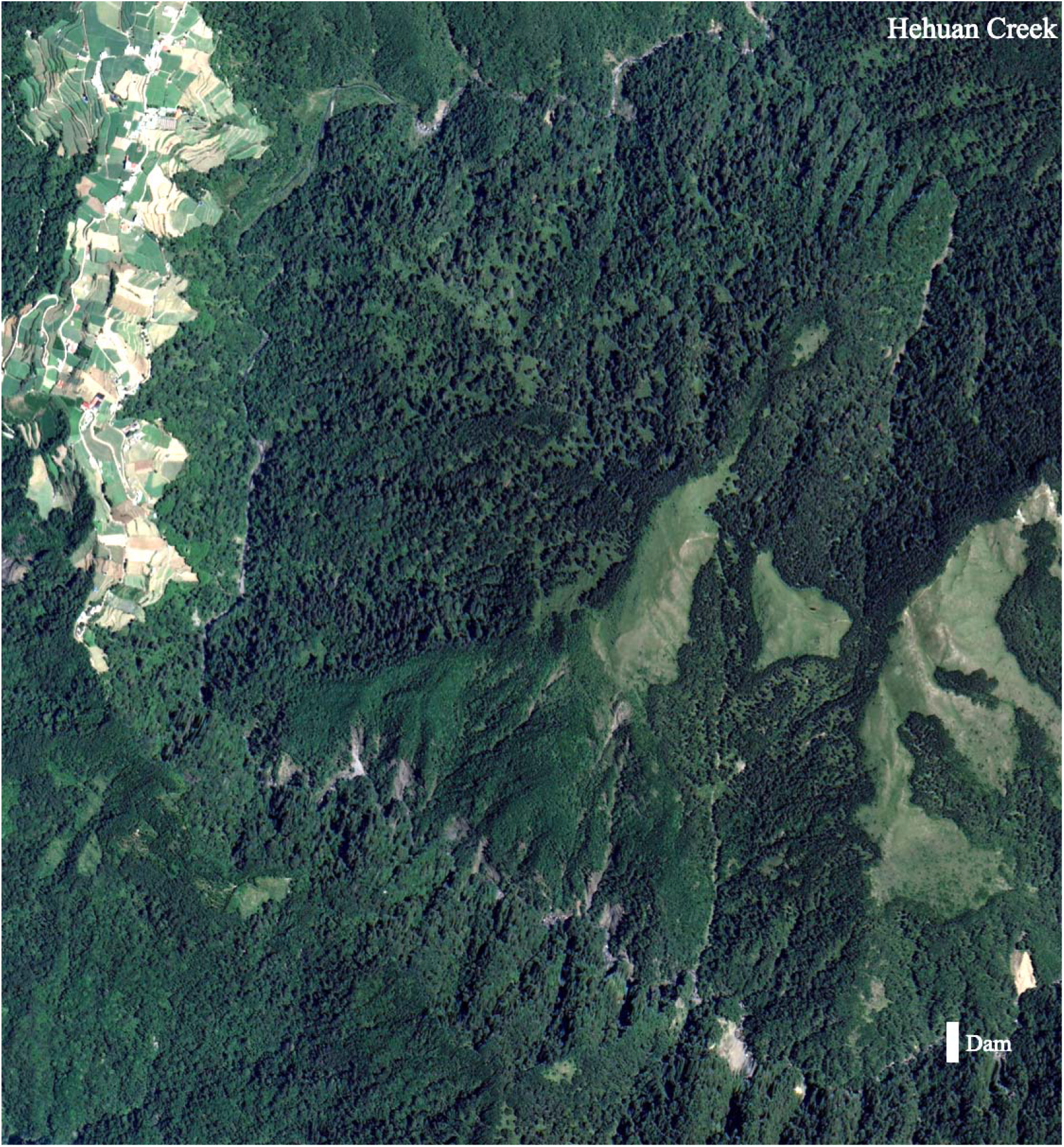
A satellite photo of the study area of Hehuan Creek. Image from Google Earth.

**Table S1.** Linear mixed model of population density and pH. Other water qualities and temperature factors were removed from the model because they were not significant.

| factors | estimate | Std. error | *t*-value | *p*-value |
| --- | --- | --- | --- | --- |
| pH | -3472.09 | 592.67 | -5.86 | 0.006 ** |

**Table. S2.** General linear model of redd site selection. Dissolved oxygen and temperature factors were removed from the model because they were not significant.

| factors | estimate | Std. error | *z*-value | *p*-value |
| --- | --- | --- | --- | --- |
| pH | -5.88 | 1.35 | -4.35 | < 0.001 *** |
| Conductivity | -100.35 | 21.58 | -4.65 | < 0.001 *** |
| Velocity | -0.11 | 0.02 | -4.64 | < 0.001 *** |

**Table S3.** AIC comparison of the linear mixed model for the first measurement data of the hatching rate.

| model | formula | AIC |
| --- | --- | --- |
| 1 | alevins = velocity + pH | 718.44 |
| 2 | alevins = velocity × Tmax + pH | 718.16 |
| 3 | alevins = velocity + pH × Tmax | 713.69 |
| 4 | alevins = velocity + pH + Tmax + DTR | 713.64 |
| 5 | alevins = velocity + pH × DTR | 709.95 |
| 6 | alevins = velocity × DTR + pH | 707.34 |

**Table S4.** The best linear mixed model for the hatching rate of the first measurement data.

| factors | estimate | Std. error | *z*-value | *p*-value |
| --- | --- | --- | --- | --- |
| velocity | -1.01 | 0.19 | -5.39 | < 0.001 *** |
| DTR | -8.12 | 2.69 | -3.02 | 0.004 ** |
| pH | -13.9 | 5.06 | -2.76 | 0.007 ** |
| velocity×DTR | 0.40 | 0.10 | 4.20 | < 0.001 *** |

**Table S5.** AIC comparison of linear mixed models for the second measurement data of hatching rate (DO means dissolved oxygen).

| model | formula | AIC |
| --- | --- | --- |
| 1 | alevins = DO + conductivity | 706.00 |
| 2 | alevins = DTR × DO + conductivity | 702.24 |
| 3 | alevins = Tmin × DO + conductivity | 700.75 |
| 4 | alevins = DO + conductivity + Tmin + DTR | 699.32 |
| 5 | alevins = DO + DTR × conductivity | 695.74 |
| 6 | alevins = DO + Tmin × conductivity | 692.08 |

**Table S6.** The best linear mixed model for the hatching rate of the second measurement data.

| factors | estimate | Std. error | z-value | *p-value* |
| --- | --- | --- | --- | --- |
| DO | -4.00 | 1.78 | -2.25 | 0.027 * |
| Tmin | -15.20 | 6.57 | -2.31 | 0.024 * |
| conductivity | -530.13 | 307.26 | -1.73 | 0.089 . |
| Tmin × conductivity | 53.43 | 31.37 | 1.70 | 0.093 . |

